# Simulating a *for-loop* in the human genome: Design and evaluation of recombinase genetic programs that count to three

**DOI:** 10.1101/2025.08.04.668540

**Authors:** George Chao, Evan Appleton, Clair S. Gutierrez, Lilia Evgeniou, Tristan Daifuku, Timothy Wannier, Esther Mintzer, George M. Church

## Abstract

Pluripotent cells specialize into numerous cell types by receiving external signals, making fate decisions, and executing differentiation functions – a paradigm similar to computer algorithms. While advances in biosensor design have enabled cells to respond to diverse stimuli, the ability to maintain a synthetic memory of the cell’s experiences that then informs its behaviors remains elusive. Here, we developed a system for cellular memory-driven behaviors by simulating a “for-loop” that counts to three using recombinase STepwise gene Expression Programs (STEPs). STEPs were genomically integrated into human cells, genotyped through targeted nanopore sequencing, and evaluated for function through changes in fluorescent reporter expression. While all STEPs were capable of heritable memory and sequential gene expression, the STEP design using tyrosine recombinases for successive excisions (TRex) significantly outperformed the others tested. We then used live cell imaging to track TRex cells as they incremented from count zero to three and observed the successive emergence of four cell states from an initially homogeneous population. The STEPs framework provides biological memory and conditional expression capabilities that, coupled with input mechanisms such as biosensors, can start to approach a programming language for biology.

## Introduction

Computers have become an indispensable and ubiquitous part of modern life due to their ability to solve countless problems. This versatility is achieved through their core design to *accept information*, *make decisions* based on input and innate conditions, then *perform functions*. Computer scientists have created a graphical representation of this concept, known as “state machines” (**Figure 1A**), to facilitate the design of complex algorithms. Biological processes, much like computer programs, can also be distilled into the same three core components. This applies across scales: from a wolf deciding to pursue prey, stem cells choosing between proliferation and differentiation, to transcription factors binding DNA targets. As the components and interactions of biological systems are characterized, it becomes possible to describe the systems as state machines and, importantly, utilize biological state machines to understand and engineer biology for therapeutic applications.

**Figure 1.**
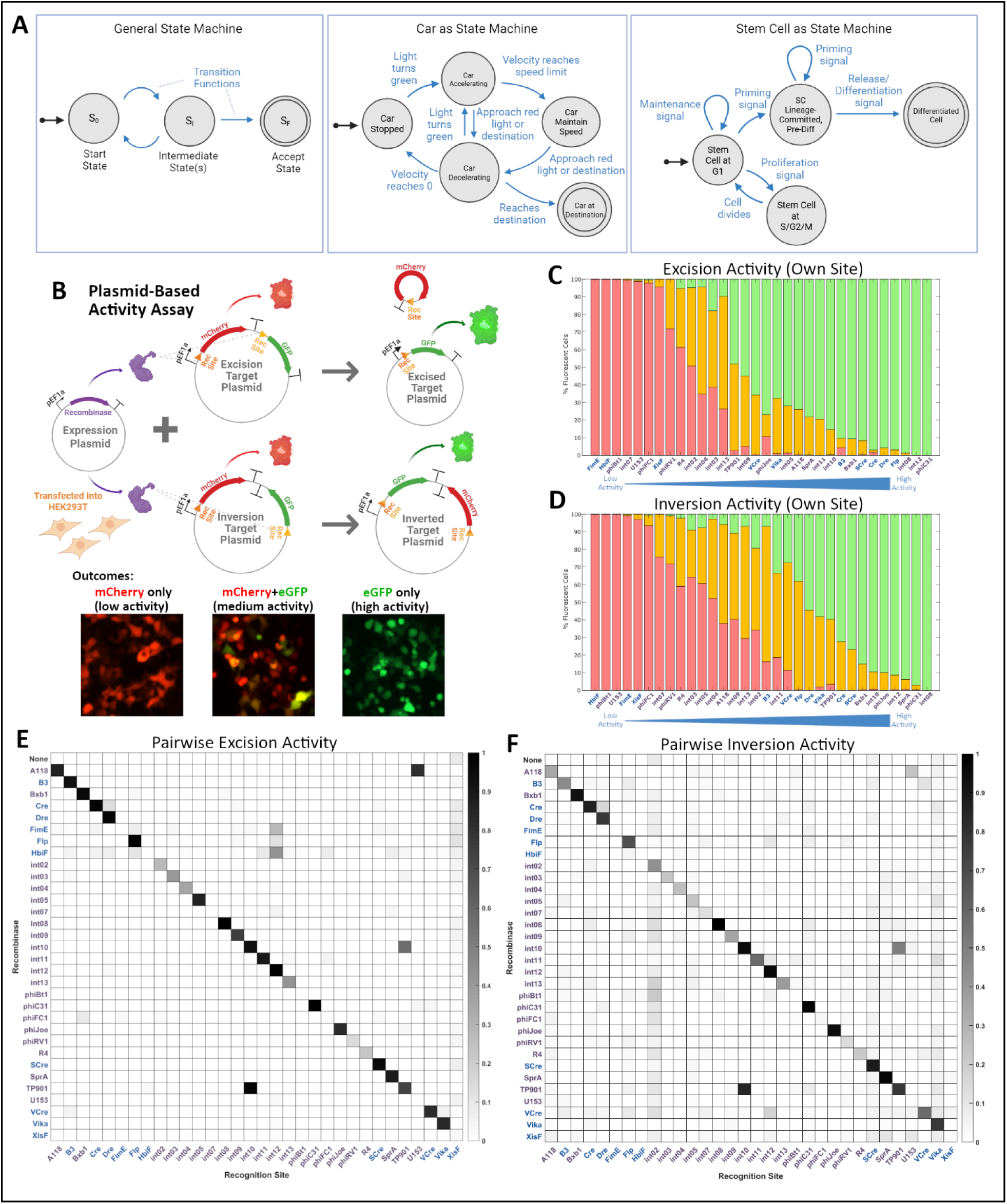
State Machine Representation of Programs and Site-Specific Recombinase Activity Assay. (**A**) From left to right: basic make-up of a state machine diagram, a state machine representation of an everyday task (controlling a car’s acceleration and deceleration), a state machine representation of a stem cell’s decision-making process. (**B**) Design of plasmid-based assay for testing recombinase activity on recognition sites (pEF1a = EF1a promoter, T symbols represent SV40 terminators). Captured at 72 hours post-transfection, recombinases with high activity on co-transfected sites present as GFP-positive cells, whereas recombinases with little activity present as mCherry-positive cells. As a dynamic process, intermediate activity can be observed in mixed populations of mCherry, GFP, and dual-positive cells. (**C, D**) Activity of recombinases on matched sites, sorted from low to high, for the excision and inversion operations, respectively. Each bar represents the mean of N=2 biological replicates. (**E, F**) Heat map of activity of all recombinases against all recognition sites for excision and inversion operations, respectively. Values represent a weighted activity score with 1 for GFP-only cells, 0.5 for dual-positive cells, and 0 for mCherry-only cells. The top row represents the plasmid-only condition for each target plasmid as a negative control. Each cell represents a mean of N=2 biological replicates.

As scientists unravel mechanisms underlying biomolecular pathways, the list of actors and their intricate relationships continues to grow. Such complex systems are challenging to harness and control, and as such, synthetic biologists are interested in creating a parallel “programming language” for biology by utilizing exogenous and often engineered genes. There has been tremendous progress in the first area of *accepting information*, with the invention of a variety of generalized biosensors utilizing RNA ^1–5^, proteolysis ^6^, GPCR signaling ^7^, and more. However, challenges remain in bridging the act of receiving information to integrating the information with innate cellular memory to *make decisions* and *perform functions*. As such, most synthetic programs today rely on a single on/off, or “Boolean,” determination. In a computer program, each Boolean decision requires a single bit of memory; by this standard, most biosensors can be considered 1-bit programs.

There have been several efforts to implement multi-bit programs in biological systems, initially in *E. coli*. One of the first is the “Repressilator,” using oscillations from an unstable inhibitory network to alternate between high and low GFP expression ^8^. Since then, other creative *E. coli*-based programs include ones that count to three ^9^, simulate complex logic ^10^, and record up to 11 bits of information ^11^, implemented as a simple genetic record without any responsive or functional elements. The last work further showcased a cascaded 3-bit program using *int02, int05, and int07*, which led to a single function change of GFP expression upon completion. Within mammalian systems, a prime example of a 1-bit program is the popular Cre-mouse system, which enables tissue-specific gene expression ^12^. Mammalian multi-bit programs have been more challenging to implement than in prokaryotes due to the higher complexity.

Notable efforts include a 3-bit program using vanillic acid-responsive mechanisms to differentiate iPSCs into beta-like cells ^13^ and a 6-bit program utilizing orthogonal tyrosine recombinases for logic gate operations ^14^. However, both rely on transient plasmid transfection, which is not suitable for long-term memory, as plasmids dilute over division cycles. Brainbow, a lineage tracing method, takes a different approach, utilizing only one recombinase (Cre) but multiple sites to combinatorially enable nine outcomes ^15^. Most current mammalian genetic programs lack one or more features critical to reproducing computer-like memory, such as memory maintenance, deterministic control, and extensibility to higher bit counts.

We hypothesize that a genome-integrated multi-recombinase genetic program (RGP) would fulfill these missing features and become a foundational tool for constructing cellular programs. Firstly, genomic integration ensures faithful proliferation of the program state and stable copy number as cells divide. Secondly, by utilizing orthogonal recombinases, each bit within the program memory is individually addressable, allowing for predictable program behavior. Thirdly, the general design principle of the programs can be learned and implemented as new orthogonal recombinases are discovered to expand the program to higher bit counts. Such a program would expand existing functionalities to be able to track multiple cell types simultaneously, turn genes on or off at different stages of development, or record occurrences of multiple intracellular or extracellular signals over time.

In this work, we describe three advances towards developing a platform for a generalized and extensible recombinase genetic program in mammalian cells. First, we tested site-specific recombinases for activity and orthogonality through a plasmid-based assay and found that of the 32 recombinases tested, 26 had detectable activity in human cells, and many were mutually orthogonal. Next, we developed a workflow for Cas9-directed integration of transgene cargos into the human genome, including a method for *de novo* discovery of on- and off-target integrations using Tn5 tagmentation and nanopore sequencing, which produced over 3,000-fold enrichment in reads of interest. We designed and tested four distinct 3-bit RGP designs using top-performing recombinases to allow cells to “count to three,” with a corresponding fluorescent reporter denoting each count. We found that, while all RGPs using top-performing recombinases were able to proceed through all three counts, the RGP utilizing tyrosine recombinases to successively excise DNA (TRex) led to considerably better performance than other designs.

Finally, we introduced the TRex program into adherent HEK293T cells and showed, using live cell microscopy, that it is possible to iterate through all three count increments within a single population without enrichment, creating a final population of cells in all four states from a homogenous progenitor population. These findings illustrate the potential for genome-integrated RGPs as a platform enabling both long-term memory and deterministic control of cell behavior, both of which are key components towards designing a generalizable framework for creating computer-like programs in mammalian cells.

## Results

### Evaluation of 32 site-specific recombinases identifies many with high activity in human cells

The primary constraint to constructing more sophisticated cellular programs is the number of “bits” that can be manipulated by the program to make and execute decisions. Within RGPs, the most straightforward way to increase the “bit” count is by incorporating additional orthogonal recombinases. Due to their versatility in gene therapy and genetic programs, there have been several efforts to mine the metagenome for site-specific recombinases. We searched the literature and curated 32 recombinases across the tyrosine recombinase (TR) and the large serine recombinase (LSR) classes ^11,16–32^. To test these recombinases’ activity in a human cell context, we designed 1-bit RGPs which expressed mCherry in its unrecombined state and GFP in its recombined state for both the excision and inversion operations and created “target” plasmids for each recombinase recognition site (e.g., *loxP*, *attB/attP,* **Figure 1B, S1**). By co-transfecting each “target” plasmid with each recombinase expression plasmid in HEK293T cells, we tested each recombinase against their own and others’ recognition sites for both operations in duplicate for a total of 4,096 transfections. On day three, we quantified the percentage of cells expressing mCherry, GFP, or both.

We found that 26 of the 32 tested recombinases have detectable activity in HEK293 cells (**Figures 1C, 1D**). Well-known recombinases such as *Cre* and *Flp* from the TR family and *Bxb1* and *phiC31* from the LSR family rank highly, as expected. Notably, several lesser-known recombinases demonstrate comparable activity to their commonly used counterparts, including *int08*, *int11*, *int12*, *A118*, *SprA*, and *phiJoe*, indicating their potential utility in gene editing and RGP applications. Our assay also reflects a key difference in the mechanism of TRs compared to LSRs. Most TRs tested act on a pair of identical sites (e.g., *loxP*), whereas LSRs bind two distinct sites (e.g., *attB/attP*) and recombine them into inactive sites (e.g., *attL/attR*). This has less impact for DNA excision, through which a free DNA molecule is formed, thus creating a thermodynamic barrier for the reverse reaction (re-integration). However, for DNA inversion, both sites remain on the same molecule post-recombination. As such, it is expected that, while LSR-catalyzed inversion reactions remain unidirectional due to the formation of inactive sites, TR-recombined sites remain active, allowing TR proteins to catalyze further DNA inversion to change the sequence back to its original state. Accordingly, performing excision with TRs led to a higher percentage of green-only cells compared to the inversion operation, where a higher percentage of cells were dual-positive. This effect was significantly less pronounced for LSRs (p-value of 0.0011, **Figure S2**).

By transforming the relative fluorescence percentages into a single weighted activity score, we visualize the pairwise activity of each recombinase against all other recognition sites for the excision and inversion operations (**Figures 1E, 1F**). Across the board, our assay generally had low noise, with only two inversion plasmids exhibiting leaky GFP expression. We attribute this to promoter-like activity from the respective recognition sites. A majority of recombinases tested have minimal activity on other recombinase sites, with the exception of *A118/U153* and *TP901/int10*. We detail our further investigations into these pairs in **Figure S3**.

### Design of 3-count RGPs using highly efficient recombinases and construction of clonal genome-integrated K562 lines

Having evaluated activity and orthogonality, we designed RGPs using top candidates. To create a universal and extensible way to track program state and translate it to selective cellular function, we designed RGPs that, with the expression of each successive recombinase, position the promoter to turn on a distinct fluorescent protein encoded in the program. In programming terms, the program function is determined by a conditional “count,” which increments with each round of recombinase expression, similar to a *for-loop*. We call these generalized “counting” RGPs “STepwise gene Expression Programs,” or STEPs. Considering constraints in the complexity of synthesis and distinguishing between downstream fluorescent reporter, the programs were designed with four distinct states, from “Count 0” to “Count 3,” with three recombinase-catalyzed transitions from mScarlet-expressing to eBFP2, iRFP670, and GFP (State Machine representations and expected genetic sequences at each step described in **Figure S4**).

We designed two configurations of STEPs – utilizing either excision or inversion operations to increment count – and utilizing either TR or LSR recognition sites, resulting in four distinct designs: TRex, SRex, TRinv, and SRinv (**Figure S1**). **Figures 2A and 2B** outline the genetic program design along with a basic state machine showing expected transitions. The excision designs are straightforward in that components are successively removed at each count, although the action of removing a section of the program is largely irreversible. Inversion-based designs are more complex to design conceptually, but all program components are retained through each count, leaving the potential of returning to earlier counts. Notably, our designs prioritized maintaining a set distance of two recognition sites between the promoter and reporter gene across all counts to minimize recognition site effects on expression ^33^. Alternative STEP designs and trade-offs are described in **Figure S5**.

**Figure 2.**
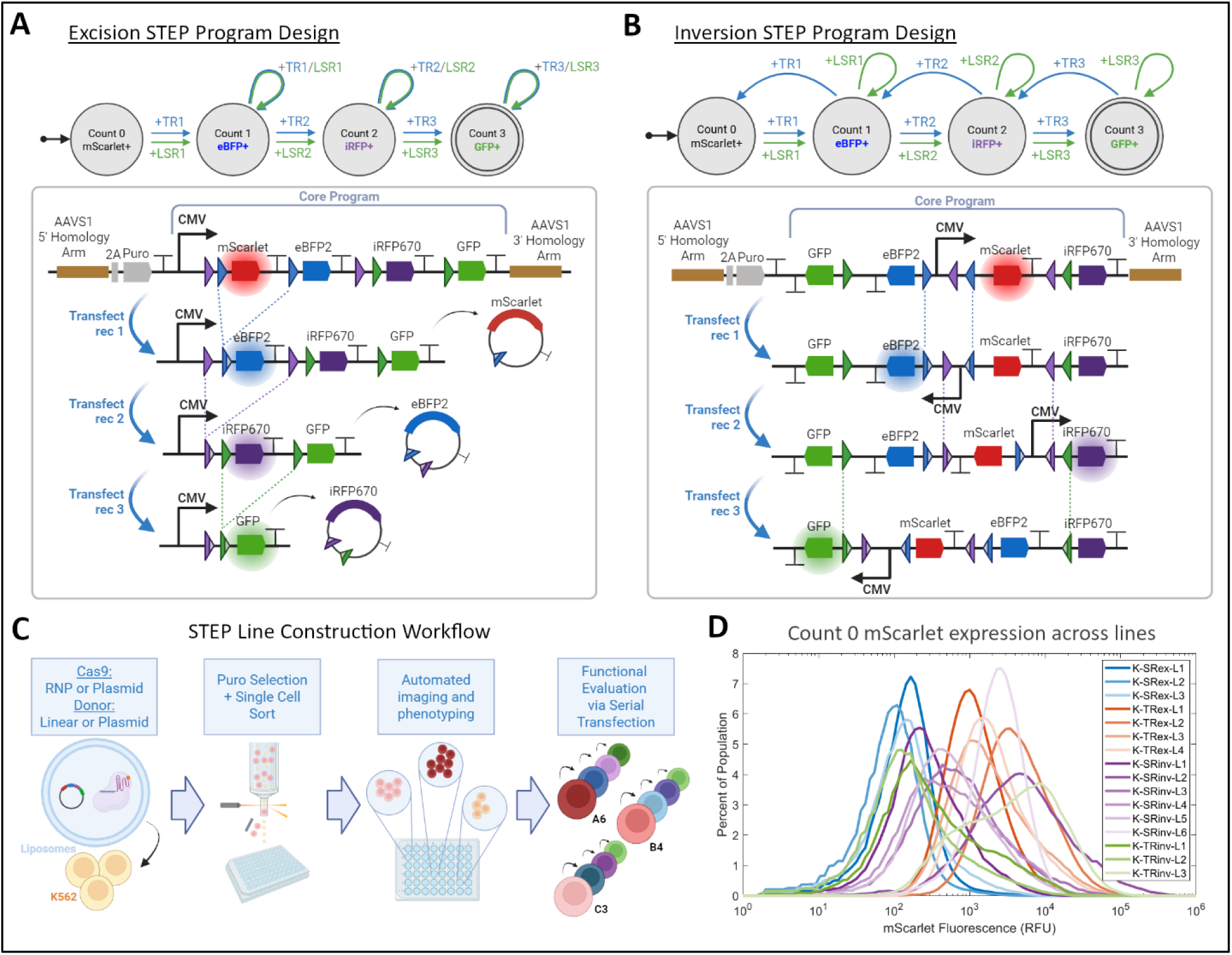
Design of recombinase programs and Cas9-guided line construction workflow. (**A-B**) General design of excision and inversion STEP programs. The excision-based program progressively cuts out the coding region and corresponding terminator, leading to the expression of the next set of genes. The inversion-based program reorganizes the position and orientation of the promoter at each count transition to induce expression of the next set of genes. A basic state machine showing the successful program sequence is shown here, and a detailed state machine of all potential outcomes is in **Figure S4**. (**C**) Workflow for targeted integration of STEP programs and clonal line construction. (**D**) Count 0 mScarlet fluorescence distribution of clonal STEP lines. SRex lines are represented by shades of blue, TRex by orange, SRinv by purple, and TRinv by green.

Next, we set out to integrate the STEP programs into the genomes of human K562 suspension cell lines. To minimize locus-specific influences on program performance and to avoid the potential for recombinases to catalyze cross-chromosomal rearrangements, we avoided random integration methods (e.g., PiggyBac or lentivirus) and utilized Cas9-directed cutting and homologous repair (Cas9-HR) to integrate our programs into the AAVS1 safe harbor site in chromosome 19 ^34^. To do this, we tested both ribonucleoprotein particle-delivered (RNP) Cas9 and plasmid-expressed Cas9 and co-delivered each alongside either linear or plasmid DNA cargoes. After delivery, cells were selected using puromycin, then single-cell sorted using mCherry to create clonal lines (**Figure 2C**). Out of 8,360 single-cell wells, 569 produced surviving clones, from which we selected 50 with high viability and mScarlet expression to test for their ability to proceed through all three counts of the program. Of these, 16, including at least three lines from each STEP design, were capable of iterating from Count 0 to Count 3.

Overall, we found that delivering Cas9 as RNPs alongside circular donors led to the lowest likelihood of undesirable outcomes (e.g., concatemerization). A detailed analysis comparing the results of each integration approach can be found in **Figure S6**).

### Development of a multiplexed and long-read method for de novo discovery of transgene integration locations

As K562 cells are triploid in chromosome 19, we expected three potential mScarlet expression levels at the Count 0 state for our cells. Unexpectedly, we observed a broad range of fluorescence across our finalized set of cell lines (**Figure 2D**). We hypothesized that this variability is due to additional off-target integrations of our program, which was confirmed by ddPCR (**Figure S7**). The most common approach to determine integration sites is to use high-coverage, short-read, whole-genome sequencing to detect genome-insert junctions. However, this approach is costly and has difficulty in uniquely mapping repeat regions. Furthermore, no single read can span from the insert across the ∼800bp homology arm, making it challenging to definitively verify even on-target integrations. Non-WGS methods exist, notably UdiTaS ^35^, Cas9-guided enrichment ^36^, and INSERT-Seq ^37^. Each method has unique limitations: short-read, low sensitivity, and requiring a ligation step (**Table S1**), respectively. The need for ligation, in particular, is undesirable for integration detection, as there is a chance of new insert-genome junction formation leading to false positives.

We therefore set out to create a method that is long-read, straightforward, affordable, and ligation-free. To do so, we combined the tagmentation and amplification components of the UdiTaS method with the ease-of-use and long-read nature of the Oxford Nanopore Rapid PCR Barcoding kit (ONT-RPB) to create a custom targeted sequencing workflow, TaP-N-Seq (TAgmentation-Pcr with Nanopore Sequencing) (**Figure 3A**). Briefly, the workflow utilizes the ONT-RPB kit-provided Tn5 transposase-transposon complex to “tagment” genomic DNA, fragmenting and adding transposon “handle” sequences randomly across the genome. Next, it uses an ONT-RPB kit-provided barcoded and chemically modified transposon primer and a user-provided primer against the sequence of interest to preferentially enrich genome-insert junction sequences through PCR. Finally, the end-labeled amplicon library is attached to the ONT-RPB kit-provided sequencing adapters in a ligase-independent manner and loaded onto an ONT sequencing flow cell. All in all, the method only requires one user-provided component of a target-specific primer and requires less than one hour of hands-on time.

**Figure 3.**
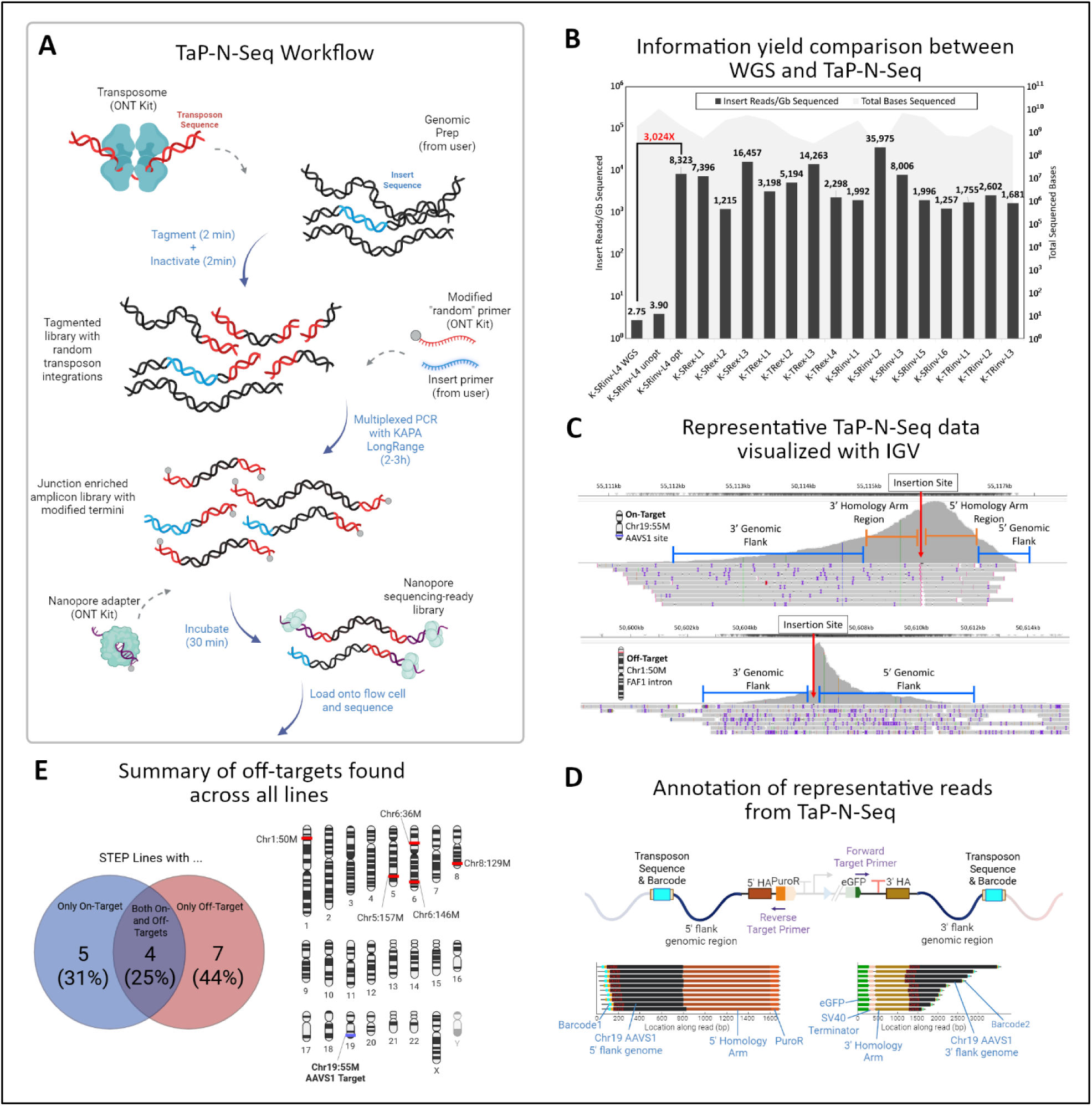
TaP-N-Seq method for *de novo* discovery of integration locations. (**A**) Details of the TaP-N-Seq method for *de novo* integration site discovery. (**B**) Comparison of informative read yield between whole genome sequencing and TaP-N-Seq of the K-SRinv-L4 line, along with the yield obtained across all 16 lines sequenced. Black bars (left y-axis) represent the number of insert reads sequenced normalized to the sequencing yield – a metric for information content. The gray area graph (right y-axis) represents the total number of bases sequenced for each TaP-N-Seq run. (**C**) Sample IGV genome-level view of TaP-N-Seq data. (**D**) Annotation of representative reads from TaP-N-Seq sequencing, demonstrating the capability for single reads to span from the insert, through the homology arm, out into the surrounding genomic region. (**E**) Summary of integration type across STEP lines and observed off-target locations.

We first tested TaP-N-Seq with minimal changes to the ONT-RPB protocol and a 1:1 ratio of the transposon primer to the target primer. This initial approach led to a modest 44% increase in the number of insert reads per gigabase sequenced (IR/GbS), an information density metric that accounts for the throughput and read length variability of nanopore sequencing.

Using a fluorescently labeled target probe, we optimized each step of the workflow to encourage preferential amplification of genome-target reads (**Figure S8**). Within the SRinv line used for optimization (K-SRinv-L4), the optimized TaP-N-Seq led to a 3,026-fold increase over WGS and a 2,134-fold increase over the unoptimized protocol (**Figure 3B**). We sequenced the remaining 15 lines with TaP-N-Seq and observed comparably high IR/GbS yields for each. With an average coverage of 1.19X, we observed an average of 14,436 insert reads per line.

We further created a computational workflow for the analysis and visualization of TaP-N-Seq data (**Figure S9**). Once mapped to the genome using conventional mapping software such as minimap2 ^38^, the alignment data can be visualized using tools such as IGV ^39^. TaP-N-Seq reads form a distinct hill-shaped profile, with tails created through random genomic fragmentation by the Tn5 transposase (**Figure 3C**). Once mapped to the genome, TaP-N-Seq reads – represented as prominent high-coverage regions – can be extracted and further annotated with insert features (**Figure 3D**).

Profiling on- and off-target transgene integration in the 16 functionally validated lines, we were surprised to find that only five (31%) had solely on-target integrations. Four (25%) had both on- and off-target integrations, and seven lines (44%) had solely off-target integrations (**Figure 3E, Table S2**). The presence of the on-target integration for each line and all detected off-target integrations were further confirmed through junction PCR and sequencing. With knowledge of flanking genomic sequences, we performed long-range PCRs spanning from the 5’ genomic flank region to the 3’ region of the insert and vice versa, allowing us to verify the entirety of each insert region (**Table S2**). We observed that on-target insertions are generally precise, with a single, unmodified copy of the desired insert and no aberration in the transition region from the homology arm into the intended genomic flank regions. However, off-target insertions are irregular, often starting from an arbitrary location within the insert, containing concatemers, and deleting sections of the surrounding genome. We further investigated the genomic context of the off-target integration sites and compared them to off-target predictions for the AAVS1-T2 gRNA from eight *in silico* and one empirical method (GUIDE-Seq)^40–46^ (**Table S3**). The Chr6:36M locus contains a clear potential off-target cut site, with only three bases distinguishing the cut site from the guide sequence. This locus was predicted by most tools tested and has also been previously reported ^47^ as an off-target for the AAVS1-T2 gRNA. Three other sites were suggested by the CasOFF tool, but only as part of an unfiltered list of over 12,000 candidates, and all other sites were not found by either *in silico* approaches or GUIDE-Seq. Our findings illustrate the prevalence of off-targets created through the Cas9-HR method, and TaP-N-Seq provides an affordable and scalable method for discovering them.

### STEP-integrated K562 cells are capable of proceeding through all three counts, with TRex significantly outperforming other designs

Having verified insert integration and quantified copy number, we next determined how well each STEP facilitates progression through all three counts. Cells were transfected with the corresponding recombinase for that count, imaged on days 3 and 6, and their fluorescence in all four channels was quantified at the population level by flow cytometry and sorted for fluorescence in the subsequent count. We found that all four STEP designs were capable of proceeding through all three transitions and, with FACS enrichment, generated pure populations of cells expressing solely the corresponding fluorescent reporter at each count (**Figures 4A, 4B, S10**), albeit with widely varying efficiencies (**Figure 4C**).

**Figure 4.**
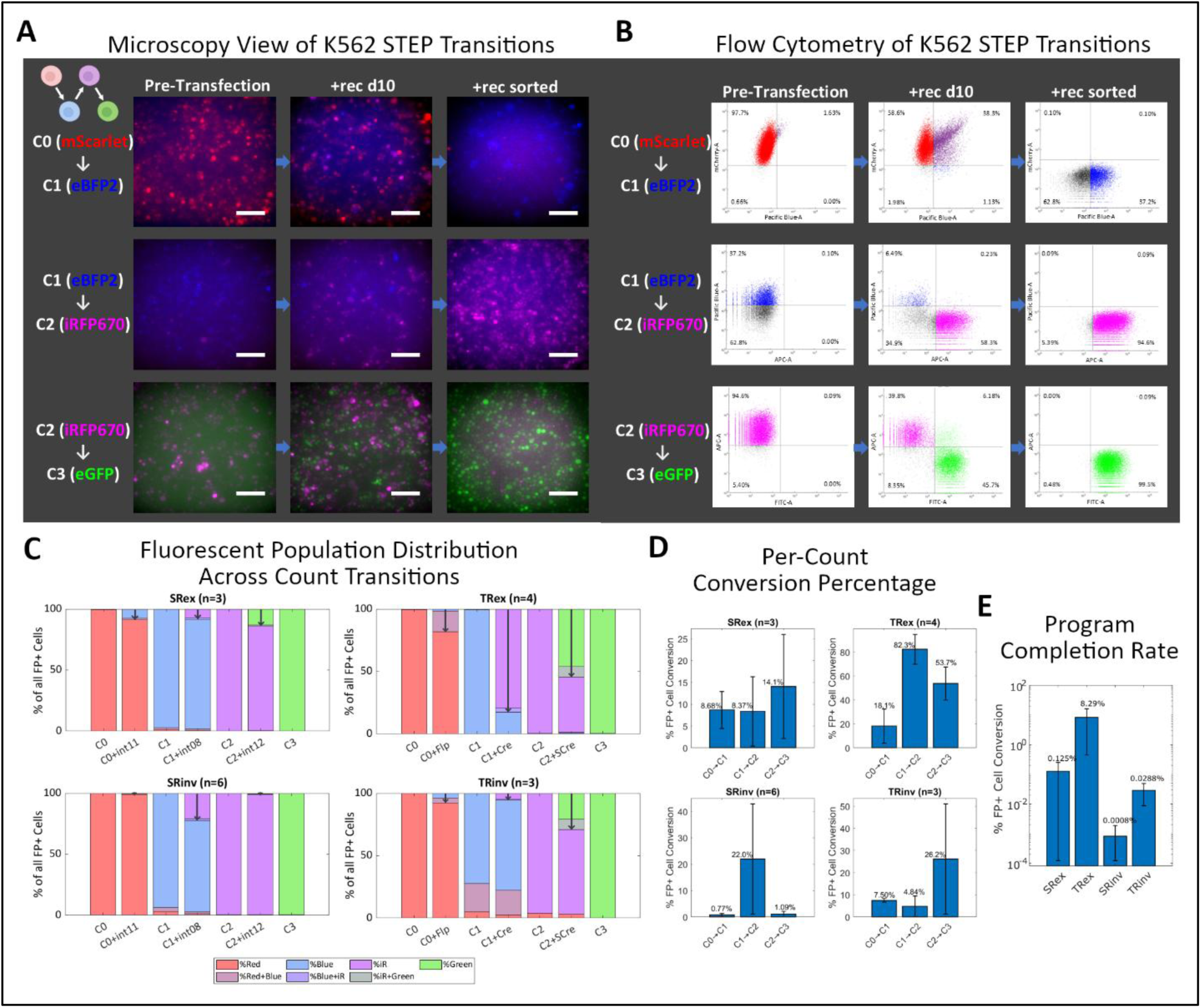
Evaluation of STEP programs in clonal K562 cells. (**A-B**) Example of STEP cell transition from C0 to C3 as observed through microscopy or flow cytometry (K-TRex-L2 line). Scale bars are 100 µm. For each row, the channel representing the starting count is on the y-axis, and the channel representing the subsequent count is on the x-axis. Cells start in the top left quadrant, transition through the top right quadrant, and end in the bottom right quadrant. This population is transfected to initiate the next count transition. (**C**) Relative abundance bar graph illustrating the change in population as cells transition from C0 to C3. Bars represent the average distribution of fluorescent cells for either a “Count X” population or a transitional “Count X+rec” population at day 10. Dark arrows represent the percentage of the population that has transitioned to the next count, which is then sorted to create a pure population of “Count X+1” cells. (**D**) Average per-count percentage conversion at day 10 for each STEP program. Note that the y-axis varies across lines. Error bars are s.d. (**E**) Average percent program completion rate for each STEP program. Y-axis is log scale, and error bars are s.d. Abbreviations: SR = Large Serine recombinase; TR = Tyrosine Recombinase; ex = excision; inv = inversion.

Across the four STEP programs, TRex had the highest conversion efficiency for all three transitions across counts (abbreviated C0-C3): 18% for C0 to C1, 82% for C1 to C2, and 54% for C2 to C3 (**Figure 4D**). The comparable LSR-based excision program, SRex, had a significantly lower conversion rate of 8.7%, 8.4%, and 14% for the respective count transitions (p-value of 0.14, <0.001, and 0.005), despite similarly high activity in earlier plasmid-based assays. In most cases, conversion efficiencies were lower for inversion programs than excision programs. As expected, catalyzing inversions with TRs sometimes led to populations that remain in a transitional state between counts (e.g., C0 to C1 transition for the TRinv line K-TRinv-L3, **Figure S10**), but generally, pure populations of the subsequent count could be derived with stringent FACS gating.

A particular metric of interest is the absolute percentage of starting cells that would successfully make all three transitions within a single, unsorted population. We calculate this “Program Completion” efficiency by multiplying the three per-step conversion percentages for each line, then averaging across lines for each program (**Figure 4E**). An ideal program would induce consistently higher conversion at each step and lead to a large percentage of cells that complete the full program. Accordingly, TRex cells demonstrated the highest program completion percentage of 8.29%, 66-fold higher than the second place SRex’s percentage of 0.13% and 288-fold higher than the inversion version’s 0.029%. Across the board, TR-based programs outperformed SR-based programs (p-values of 0.065 for excision and 0.069 for inversion), and excision-based programs outperformed inversion-based programs (p-values of 0.063 and 0.12).

### Dynamics of complete STEP program execution observed through live cell imaging of HEK293T TRex cells

For STEPs to be portable and versatile, they would ideally function across multiple cell types and generate a substantial subpopulation of “successful” cells without the need for enrichment. As TRex exhibited the best performance of the four designs, we selected it for further testing and integrated it into the genome of adherent HEK293T cells using the same Cas9-HR strategy as above to create clonal lines. We then transitioned the cells from C0 to C3 without enrichment, tracking the dynamic changes in the four fluorescent reporters using flow cytometry (**Figure S11**) and live cell imaging. The flow cytometry data showed that the TRex program behaved similarly in HEK293Ts as it previously did in K562 cells, with correlated conversion efficiencies at each transition (R^2^ = 0.80, **Figure S11**).

The live cell imaging experiment, spanning approximately 30 days and presented both as a time-lapse video (**Movie S1**) and a panel (**Figure 5**), provided striking visual confirmation of the program’s behavior. With each transition, a new population of cells emerged that expressed the fluorescent reporter for the subsequent count, leading the population to grow increasingly heterogeneous through the experimental period. The imaging data also captured the kinetics of gene expression following count transitions: each subsequent fluorescent reporter became visible around 48 hours post-transfection and continued to rise over eight days. Throughout the experiment, HEK293T TRex cells maintained strong reporter expression while continuing to divide and proliferate, indicating that the program operated in parallel to the cells’ native functions. Together, these results demonstrate that TRex-integrated cells remain viable and functional across all counts, highlighting their suitability for use in complex multicellular systems.

**Figure 5.**
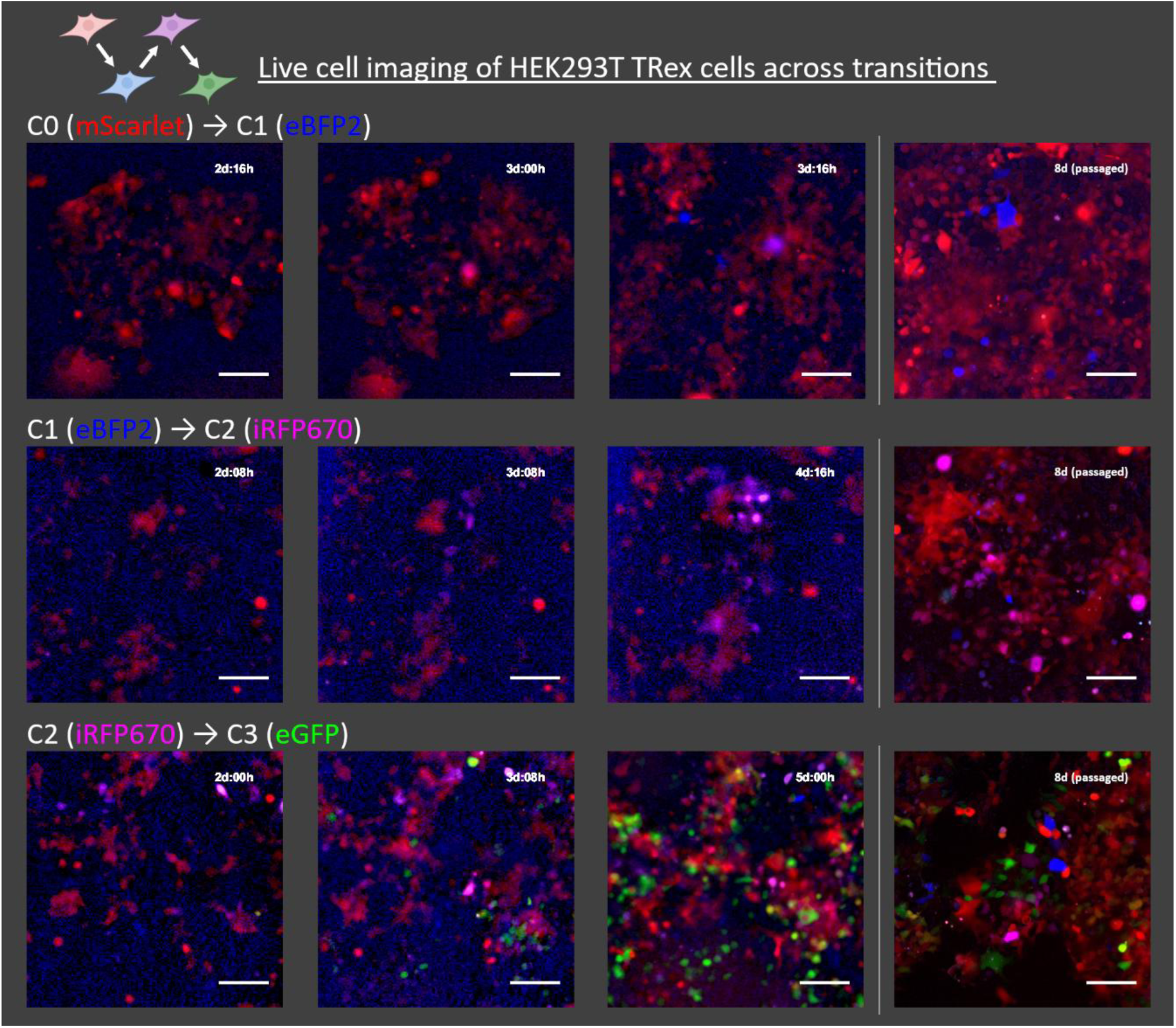
Live cell imaging of a single HEK293T TRex population from C0 to C3. Panels represent selected, cropped images from live-cell imaging of a HEK293T TRex cell line as it is successively transfected with recombinases. The population was not enriched at any point, demonstrating the emergence of four distinct and stable cell populations stemming from a single, homogenous starting population. Images that demonstrate progressive change in the subsequent count’s fluorescence are shown. Live cell images were taken using a 10X objective using a 2x2 stitched conformation (cropped in this view). Cells are passaged at 5 days post-transfection and imaged at day 8 using a 20X objective for higher resolution. All images show all four channels (mCherry, DAPI, Cy5, and FITC) merged. **Movie S1** shows the full, uncropped video across all transitions. Scale bars are 100 µm.

## Discussion

In this work, we outlined an end-to-end strategy for designing, delivering, characterizing, and validating genome-integrated recombinase genetic programs to retain cellular memory and perform output functions in human cells. We assayed the activity and orthogonality of 32 recombinases and found several with high activity. The most efficient subsets were selected to design four recombinase programs capable of counting and directing corresponding gene expression (STEPs) and were integrated into K562 genomes through Cas9-HR. Upon evaluating STEP cells, all four programs completed sequential transitions from Count 0 to 3, although TR-based STEPs outperformed LSR-based STEPs, with TRex scoring the highest using an integrated metric for program performance. Finally, we tested TRex in HEK293T cells, demonstrating its portability across cell lines and the ability to iterate through all three counts within a single TRex cell population without enrichment.

Notably, we observed a disparity in recombination efficiency between our plasmid-based assay and the performance of the genome-integrated STEP program, with the LSRs deviating the most. Other genomic applications of LSRs have also found similar, lower-than-expected efficiencies, such as integration into landing pads ^48–50^. A potential contributor to this difference may be the dense, heavily regulated genomic context, within which recognition sites are more difficult for recombinase proteins to access and manipulate compared to the less constrained and more abundant plasmids. Interestingly, this effect is more attenuated in TRs, particularly in the case of Cre. Conceptually, LSRs, in principle, would be better suited for simulating the “toggle” characteristic (turn on/off) of computer “bits” due to their unidirectional recombination reaction and the potential to couple with a cofactor ^51^ for the inverse reaction. For now, the comparatively low genomic editing efficiencies of LSRs hinder their current utility within STEP programs but also point to potential mechanistic factors that, with some engineering, may allow them to achieve higher efficiency (an example being eeBxb1 ^50^).

To genotype STEP lines, we developed a method for *de novo* discovery of transgene integration site, called TaP-N-Seq. Optimized TaP-N-Seq enriched the number of informative reads by over 3,000-fold over WGS. Using this method, we found several off-target integrations in our clonal STEP lines, with many of these locations not predicted by off-target prediction tools. Through this process, it became apparent that the *de novo* identification of transgene integration loci remains challenging and expensive. As such, high-quality genotyping remains rare despite the widespread usage of Cas9-HR for targeted integration and random integration methods such as PiggyBac and lentivirus. Additionally, as genetically engineered autologous cells become FDA-approved, such as Kymriah for blood cancer ^52^, Casgevy ^53^ and Lyfgenia ^54^ for Sickle Cell, and Skysona for CALD ^55^, it becomes even more pressing to have robust and affordable techniques available to encourage cell therapy manufacturers to determine integration loci, lest their patients end up trading one disease for another.

Our top-performing STEP, TRex, has the ability to simulate a *for-loop* by counting recombinase expression events and expressing effector genes in response, making it a powerful tool for reproducing temporal phenomena across several fields of biology. One compelling application is to utilize TRex to record cellular events, such as expanding upon existing recombinase-based lineage tracking methods ^56^. Along the theme of developmental biology, it should be possible to leverage biosensors for cell type-specific signals to drive TRex progression ^1–5^, which in turn can direct the expression of sets of transcription factors to catalyze desired cell fate decisions. In immune effector cells (e.g., cytotoxic T cells, NK cells, etc.), synthetic cell surface receptors can transduce spatial context information to drive TRex progression ^6^, guiding cells to migrate to specific tissues before triggering target cell killing, cytokine secretion, or other immune responses upon antigen detection. The modular design of TRex allows it to be adapted, with some creativity, for use in a broad range of applications.

Within its potential applications, STEPs fill the role of cellular memory for *making decisions* and *performing corresponding functions*. By integrating STEPs with methods for *accepting information*, the resulting genetic program would be capable of mirroring the logic of computer algorithms and thus enabling the composition of higher-complexity cellular behaviors. With continued refinements to improve efficiency and bit count, the STEP system could evolve into a comprehensive instruction set for programming cellular behaviors. More broadly, as technological advances give researchers increasingly precise control over biological functions in human cells, it becomes ever more feasible to engineer next-generation cell and tissue therapies to address some of medicine’s toughest challenges.

## Resource Availability

All raw sequencing data from TaP-N-Seq as well as MATLAB code for read annotation will be made available upon publication. All recombinase expression, excision, and inversion plasmids, as well as plasmids for all four STEP programs, will be deposited to Addgene after publication.

## Acknowledgments

We thank Integrated DNA Technologies, particularly Gavin Kurgan, Kyle Kinney, Shane Lennon, and Mark Behlke, for helpful discussions on *in silico* gRNA off-target predictions and for providing GUIDE-seq data for the AAVS1-T2 gRNA.

The plasmid used for plasmid-based Cas9 and AAVS1-T2 gRNA expression was a gift from Masato Kanemaki (Addgene 72833).

We are grateful to Dr. Bridget Baumgartner, Dr. Justin Gallivan, Dr. Jesse Dill, and Dr. Joseph Pomerening for helping to fund and shape the direction of our work.

We also thank the HMS Immunology Flow Cytometry facility, particularly Chad Araneo, Jeff Nelson, and Meegan Sleeper, for training and maintenance of the FACS machines.

We thank Dr. Paula Montero Llopis and Ryan Stephansky from the HMS MicRoN core facility for their assistance in microscopy techniques and maintenance of microscopes. We also thank Josh Rosenberg with Nikon for his responsive and expert assistance in procuring and supporting our Nikon microscope and Geoff Guimaraes from Zeiss for helping to source, set up, and support the live cell imaging system.

We are extremely grateful to Tiffany Dill and Matthew Butnaru for their help in editing the manuscript.

Thank you to Emma Taddeo and Esme Philips for their help with the lab’s administrative work, Jen Shay for lab logistics support, and Nicole D’Aleo for her assistance with budgeting for the project and grant administration.

Plasmid design and map image generation were done in A Plasmid Editor (ApE). A majority of schematic figures made in this work were created with BioRender.

## Funding

G.C., E.A., C.G., T.D., and T.W. were supported by the DARPA Engineered Living Materials program under contract W911NF-17-2-0079. G.C. was also supported by the NHGRI Centers of Excellence in Genomic Science RM1HG008525. T.W. was also supported by the DoE award DE-FG02-02ER63445. E.M. was supported by an NIH T32 grant for the Program in Genetics and Genomics (5T32GM141745-03).

## Author Contributions

Conceptualization, G.C., T.W., E.A., and G.M.C.; Methodology, G.C., E.M., L.E., C.S.G., E.A., and T.D.; Investigation, G.C., C.S.G., L.E., T.D., E.A., and E.M.; Analysis/Software, G.C. and L.E.; Writing – Original Draft, G.C.; Writing – Review & Editing, G.C., L.E., E.M., C.S.G., and G.M.C.; Visualization, G.C., L.E., and E.M.; Supervision, G.C., E.A., and G.M.C., Funding Acquisition, G.C., E.A., T.W., and G.M.C., Resources, G.M.C.

## Declaration of Interests

Full disclosure for G.M.C. is available at http://arep.med.harvard.edu/gmc/tech.html. All other authors declare no competing interests.

## Supplemental Information

Supplemental Information will be available in publication.

## Materials and Methods

### Plasmid Construction

Recombinase sequences were curated from the literature. The sequences were codon-optimized for mammalian expression and synthesized through GenScript into standard plasmid backbones. Recombinase coding regions were then PCR amplified using NEB Q5 polymerase (Cat# M0491L) and cloned via Gibson Assembly (NEB Cat#2611) into a standard expression vector with an upstream EF1a promoter and downstream BGH terminator to create recombinase expression vectors. For excision and inversion vectors, a template plasmid was first built for either the excision or the inversion orientation. Sites were then exchanged using the Agilent Multi Site-Directed Mutagenesis Kit (Cat# 200514) and single oligonucleotides encoding the new site with ∼15 bases flanking each end.

STEP program plasmids were first designed in ApE. The sequences were then directly synthesized into a custom vector using Thermo Fisher’s GeneArt Gene Synthesis service.

All plasmids for transfection were maxiprepped using the ZymoPure II Plasmid Maxiprep Kit (Cat# D4203), including the endotoxin removal step.

### General Cell Culture and Plasmid Transfections

HEK293T cells were cultured to ∼70% confluency. Transfections were performed using Lipofectamine 2000 (Cat# 11668019) following the recommended protocol, using 1.6ug of DNA with 25µL of reagent for 12-well plates and scaling based on plate surface area for other well formats.

K562 cells were cultured to near confluency based on cell count. Reverse transfections were performed in 12-well plates, where the lipofectamine-DNA solution was prepared in the same way as for HEK293T transfections but added to the bottom of empty wells and allowed to coat the entire surface. K562 cells were then counted and a cell count number of ∼25% of confluency (1.25x10^5^ for 12-well) was spun down and resuspended in culture media (1.5mL for 12-well) and added to the reagent-coated wells.

Both cell types were cultured in DMEM (high glucose, +GlutaMax, ThermoFisher Cat# 10566016) with 10% FBS (Corning Cat# 35-010-CV) and 1% Pennicillin-Streptocycin (ThermoFisher Cat# 15140122).

### Pairwise Recombinase-Recognition Site Activity Assay

Due to the quantity of transfections, 48-well plates were used to reduce reagent use. Transfections largely follow the protocol listed above. Since this assay requires the co-transfection of both an expression vector and either an excision or inversion target vector, the amount of each plasmid added was halved (0.2ug each for 48-well). Both vectors were added to Opti-MEM (ThermoFisher Cat# 31985062) during the pre-mixing step of reagent preparation. For no-recombinase control conditions, a non-expressing pUC19 vector (NEB Cat# N3041S) was added instead.

After adding the lipofectamine-DNA reagent, cells were incubated for 72 hours. They were then trypsinized (ThermoFisher Cat# 12604013) for 5 minutes, mechanically homogenized by gentle pipetting, then 10µL of the cell/trypsin solution was directly added to a chamber of automated cell counter slide (VWR Cat# 10027-446). The cells were counted using a ThermoFisher Countess II automated cell counter with EVOS fluorescence light cubes for Texas Red (ThermoFisher Cat# AMEP4955) and GFP (ThermoFisher Cat# AMEP4951).

Raw cell counts for each channel were normalized by the total number of fluorescent cells counted, then represented either as a stacked bar graph of GFP+ (green), dual+ (yellow), or mCherry+ (red) or as a single weighted value. Weighted activity was calculated using the formula:

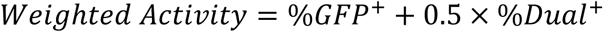

Each pairwise transfection was performed in duplicate, and weighted activity scores were compared. If the difference between the two measurements was greater than 0.25, the co-transfection was repeated a third time as a tie-breaker.

### Delivery of Cas9, gRNA, and STEP program construct for Cas9-HR

Four approaches were tried to achieve Cas9-HR-mediate integration of STEP program constructs into the AAVS1 safe harbor site. All approaches utilized the AAVS1-T2 gRNA (*57*) for targeting.

We introduced Cas9 and gRNA via either plasmid transfection or RNP delivery. For the transfection approach, we co-transfected a plasmid with a CAG promoter-driven SpCas9 with an N-terminal SV40 NLS sequence along with the AAVS1-T2 gRNA under a U6 promoter (Addgene 72833). This was done in a 1:2 (cas9:donor) ratio using Lipofectamine 2000. For the RNP approach, gRNA was chemically synthesized through ThermoFisher’s TrueGuide Synthetic gRNA service, which includes both 2’ O-Methyl and phosphothioated bonds on terminal nucleotides. This was resuspended in TE buffer (MilliporeSigma Cat# 93302) to create a 10X stock (100uM). The stock was then diluted to 1X with ultrapure, RNAase-free water (ThermoFisher Cat# AM9915G) prior to use. For Cas9, the ThermoFisher TruCut Cas9 Protein v2 (wildtype SpCas9 with 2x NLS tags) was used (ThermoFisher Cat# A36498). RNP formation largely follows the Lipofectamine CRISPRMax protocol (ThermoFisher Cat# CMAX00001). First, 50µL of Opti-MEM, 2.5µL of the stock Cas9 protein reagent, 1.5µL of 1X gRNA solution, and 5µL of Cas9 Plus Reagent were mixed and allowed to incubate for 5 minutes for RNP formation. 1ug of donor DNA was added at this point and mixed by gentle pipetting. Per protocol, 50µL of Opti-MEM and 3µL of Lipofectamine CRISPRMax reagent are mixed in a separate tube, incubated for 1 minute, then combined with the RNP+donor solution. The combined solution is mixed by gentle pipetting and incubated for 15 minutes at RT, then the entire solution is added to one well of a 12-well plate, following standard transfection protocols described above.

Donor DNA was co-delivered either as plasmid DNA containing the full plasmid backbone or as linearized DNA, which is the plasmid DNA digested with restriction enzymes PciI (NEB Cat# R0655S) and SspI (NEB Cat# R0132, the non-HF version with higher activity in NEBuffer 3.1) and gel-extracted.

### Flow Cytometry

All flow cytometry and sorting of cells were performed using a FACS Aria II from the Harvard Medical School Immunology FACS Core using a 100um nozzle. In addition to standard FSC and SSC, four channels were captured: BFP/Pacific Blue (405nm laser, 450/50 filter), GFP/FITC (488nm laser, 530/30 filter), mCherry/mScarlet (594nm laser, 610/20 filter), and iRFP/APC (633nm laser, 675/20 filter). In general, cells were homogenized, spun down, and resuspended in culture media, then passed through a mesh cell strainer into 5mL sort tubes (Corning Cat# 352235). Cells were sorted into 500µL of culture media, after which they were spun down and resuspended in the appropriate volume for culture.

### Selection of Clonal STEP Lines

After Cas9, gRNA, and STEP program delivery as described above, cells were incubated to allow for integration and expression of puromycin resistance. After 72 hours, cells were passaged into a 6-well plate containing culture media with puromycin (ThermoFisher Cat# A1113803). For HEK293Ts, a final puromycin concentration of 2ug/mL was used, whereas K562 cells were more sensitive, and a concentration of 0.5ug/mL was used. Cells were incubated with puromycin for 3 days, after which they were passaged into a fresh 6-well plate with standard culture media without puromycin and allowed to recover. Once clones of ∼20 cells were visible (for either cell type), the cells were gated for the top 10% of mScarlet expression, and single cells were sorted into 150µL of culture media in 96-well plates. Post-sorting, the 96-well plates are spun down to precipitate cells. Plates are then incubated for 15 days, at which point clones will be visible in wells with surviving cells. These wells are marked, and at ∼20 days post-sort, they are passaged into 24-well plates.

At the 24-well stage, clonal cell populations are phenotyped for mScarlet expression using a Nikon Ti2 inverted microscope system (more details on configuration in the Microscopy section). Cells are marked using the multi-point feature and stitching a 2-by-2 field of view using a 20X/0.5N.A. objective using a fixed capture setting. Images are captured and exported as 12-bit TIFF files, which were then segmented and analyzed via CellProfiler (*58*) for mean fluorescence intensity. Each well was also manually scored for viability. Top candidates for fluorescence and viability for each genotype were then selected for functional characterization by sequential recombinase expression.

### Functional characterization and analysis of STEP Lines’ Per-Step Conversion Efficiency

For STEP K562 cell lines, cells at each count were transfected in 12-well plates as per above with the corresponding recombinase for the program type. Cells were imaged at day 3 and day 6 post-transfection, then passaged into a 6-well plate. On day 10 post-transfection, they were imaged again, then measured by flow cytometry, and sorted for cells either solely positive for the fluorescent reporter representing the next count or for both single- and dual-positive cells, depending on conversion efficiency. These sorted cells were then sequentially expanded and enriched until either less than 1% of cells from the previous count remained or no further enrichment of cells in the new count. This entire process is then repeated until a pure population of GFP+ cells, representing Count 3, is observed.

The composition of fluorescent cells at each count is characterized through an automated flow cytometry analysis pipeline implemented in MATLAB. In short, “live cell” and “cell debris” gates are detected and drawn based on a peak-finding algorithm, then the “live cell” population is gated utilizing a combination of a density cutoff and an FSC/SSC distance threshold from the peak. Single cell population gates are selected for FSC and SSC based on a 99th percentile threshold. For each run, there is a wildtype K562 sample, which serves as its unstained negative control. The pipeline utilizes the unstained control to set quartile gates for each pair of consecutive count reporters and to derive a geometric mean used for normalization. Percentages are normalized to include only cells above negative control thresholds. Fluorescence is generally only reported for “pure” populations (accepted “count” populations based on previously described criteria) and represents the geometric mean of the entire live, singlet population as a ratio to the date-matched negative control geometric mean in that channel.

### Digital droplet PCR characterization of STEP integration copy number

To prepare the PCR reaction mix, 20ng of genomic DNA (DNeasy Blood and Tissue Kit, Qiagen Cat# 69504) with 10µL of ddPCR Supermix (BioRad Cat# 1863024), 1µL of reference probe mix (HEX dye), 1µL of a custom probe mix (FAM dye, synthesized by BioRad), 1µL of EcoRI-HF, and water to reach a 20µL reaction volume. Droplets were generated using the QX100 Droplet Generator following standard protocol and using BioRad consumables. Droplets were then carefully transferred to an Eppendorf semi-skirted 96-well PCR plate (Eppendorf Cat# 951020427) and covered using a foil heat seal (BioRad Cat# 1814040). The plate was thermocycled using a standard thermocycler following BioRad protocol. The reaction is then immediately measured using a BioRad QX100 droplet reader.

For each line, we tested mScarlet copy number against three loci: NSUN3 (chr3 centromeric, BioRad assay# dHsaCP2506682), RPP30 (chr10, BioRad assay# dHsaCP2500350), and GINS1 (chr20 centromeric, BioRad assay# dHSACP2506711), all of which have two copies in the K562 genome. Reported copy numbers are an average of the three assays, which generally closely align.

### Fluorescent Amplicon Enrichment Assay for TaP-N-Seq optimization

To optimize the TaP-N-Seq protocol for targeted read enrichment without needing to expend flow cells, we devised a gel quantification-based approach that only expends the FRM kit reagent. An unmodified version of the RLB primer from the SQK-RPB114 kit was synthesized, which is composed of the 5’ flank, barcode01 sequence, and the 3’ flank. The 3’ flank contains the annealing sequence for the transposon sequence randomly integrated during tagmentation. We also synthesized a modified target probe against the puromycin resistance gene with a 5’ Alexa647 label. The Alexa647 probe is spectrally distinct from our DNA dye (Sybr Gold), allowing gel imagers capable of capturing the Cy5 channel to independently image and measure total DNA and probe-incorporated DNA. Tagmentation and cycling conditions were optimized as described in **fig. S8**. The PCR reactions were then cleaned up using the Qiaquick PCR Purification Kit (Qiagen Cat# 28104) and ran on a pre-cast 1% agarose gel containing Sybr Gold (ThermoFisher Cat# G402021) for 8 minutes. The gel is then imaged using a BioRad ChemiDoc MP using standard gel conditions (trans-blue 450-490nm excitation + 590/110 broad spectrum emission filter) and Cy5 condition (Epi-red 625-650nm excitation + 675-725nm emission filter). Gels were analyzed using the BioRad ImageLab software. After background subtraction, enrichment was calculated as a ratio of the integrated Alexa675 fluorescence across the lane (amplicons incorporating the target primer) to Sybr Gold fluorescence (both random and targeted amplicons).

### TaP-N-Seq Sequencing Workflow

TaP-N-Seq is a highly optimized workflow. Genomic DNA should be prepared with the Qiagen DNeasy Blood and Tissue Kit (Qiagen Cat# 69504). High molecular weight preparations do not benefit output read lengths. Output read lengths are primarily bottlenecked by the PCR step, and longer input genomic fragments will likely be detrimental to junction read enrichment. Genomic DNA concentrations should be measured using a dye-based assay (such as Qubit Broad Range dsDNA Assay, ThermoFisher Cat# Q32853). Volumes, incubation temperature, incubation time, and handling conditions should be carefully adhered to for optimal results. The below protocol is designed for sequencing both 5’ and 3’ flanking genomic regions in a single reaction and can be adapted for single-flank or multi-site sequencing depending on coverage needs. All reagents other than the target primer can be found in Oxford Nanopore’s Rapid PCR Barcoding Kit (Oxford Nanopore Cat# SQK-RPB114.24).

Tagmenting genomic DNA with handle sequences: Prior to starting the protocol, set a PCR machine to 80°C. In a single tube, combine 39µL of genomic DNA containing 1.04ug of DNA with 13µL of FRM and immediate vortex. Incubate at RT for 2 minutes. During the incubation, aliquot 4µL of the reaction into each tube of two 6-tube strips of PCR tubes. Inactivate at 80°C for 2 minutes, then transfer to ice to cool.

Enriching genome-target DNA fraction and addition of end-modifications for rapid attachment: For each set of six tubes, add 4µL of tagmented DNA, 25µL of NEB LongAmp Taq MasterMix (NEB Cat# M0287L), 20µL water, 0.2µL of a selected RLB barcode primer, and 0.8µL of your custom target primer. Note that designing a specific, high-performance target primer is critical for successful junction read enrichment. Run a two-step PCR following: Initial Denaturation of 3 min at 95°C, X cycles of 15s denaturation at 95°C and combined anneal and extension step for 6 min 15s at 65°C, followed by a final extension time of 6 min 15s at 65°C. The number of cycles needs to be empirically determined for different target primers. We recommend initially testing both 25 and 30 cycles, then running the PCR reactions out on a gel. The goal is to achieve the highest number of cycles while maintaining a smooth smear and avoiding bands from forming.

DNA purification and attachment to pore adapters: Add 4µL EDTA to each 50µL reaction. Combine each set of 6 50µL PCR reactions into a single tube, then perform a standard PCR clean-up using the Qiaquick PCR Purification Kit (Qiagen Cat# 28104), eluting into 35µL of water. A second round of clean-up and concentration is then performed using standard AmPure XP beads at 0.6X (included in ONT kit), eluting into 15µL of ONT Elution Buffer. We do QC at this point using 1µL each for Nanodrop for purity, Qubit for concentration, and gel for size. In a separate tube, we add 0.6µL of concentrated adapter (RA) into 1.4µL of Adapter Buffer (ADB), mix by pipetting, and add 1µL of diluted adapter to 10µL of the cleaned PCR reaction. We incubate this mixture for 30 minutes.

Loading and sequencing: Following the adapter attachment, the flow cell loading and sequencing workflow follows the standard protocol from Oxford Nanopore.

### TaP-N-Seq Analysis Workflow

Nanopore sequencing generates a unique type of long-read and error-prone data (∼92-98% single molecule per-base accuracy). As such, we combined existing packages with a custom analysis pipeline in MATLAB.

FASTQ basecalling and large feature mapping: For basecalling, we use the Guppy/Dorado software as part of Oxford Nanopore’s MinKNOW software package. Through the GPU-based live basecalling feature, it is possible to get an idea of the results of the run within the first five minutes of sequencing. The output FASTA data is then mapped separately to a GRCh38 reference genome file that is stripped of alternative, unmapped, and mitochondrial sequences to generate a genome-specific SAM file using *minimap2* (*36*). The alignment is also performed against “large” features (defined as features >100bp), creating a separate feature-specific SAM file. This process creates two files that contain redundant read sequence information but simplifies downstream analysis.

Mapping of small features, barcode calling, and read-level annotation of features: The two SAM files are jointly integrated into a single data structure within MATLAB using matching Read IDs, and only primary alignments are accepted. All reads are then aligned to the expected barcode sequences using local alignment (Smith-Waterman). By normalizing read scores to the barcode length, a normalized score is derived in the range of 0-1.4, within which a threshold is set for barcode calling. At this point, hits against reference genomic sequences (Actin, RPP30) are tallied for QC purposes. Reads that mapped to large synthetic features are then extracted and aligned against short synthetic features (<100bp), and all features are annotated as graphical representations and GenBank files.

Visualization of TaP-N-Seq-enriched regions: From the above workflow, we detect reads with genomic and insert reads. This data allows for the detection and visualization of integration locations. The genomic-specific SAM file can be sorted and indexed using *igvtools* and imported into IGV (*37*) for visualization.

### In silico gRNA off-target prediction

The AAVS1-T2 gRNA sequence was inputted into eight separate tools (CasOFF, CCTop, CHOPCHOP, COSMID, CRISPOR, OffSpotter, IDT’s gRNA design tool, and Synthego’s gRNA design tool). Where possible, standard settings were used. In some cases, certain tools allow users to generate an extensive list of targets (over 10,000 for CasOFF), in which case the lists are treated separately. The generated lists are imported into MATLAB, and the list of TaP-N-Seq off targets observed was checked against the list, using a distance threshold of 100kb.

### GUIDE-Seq

AAVS1-T2 off-targets were also assessed using the empirical GUIDE-seq method (*44*). In brief, HEK293 cells that constitutively express *S.p.* Cas9 nuclease (HEK293-Cas9; ATCC: CRL-1573Cas9) were used as a model system for nominating off-targets. Alt-R gRNA complexes (IDT) were formed by combining Alt-R tracrRNA and Alt-R crRNA XT at a 1:1 molar ratio.

The Alt-R gRNA complexes, along with the GUIDE-seq double-stranded oligodeoxynucleotide (dsODN), were delivered by nucleofection using the Lonza 4D-Nucleofector 96-well unit. For each nucleofection, 2.0 x 10^5^ HEK293-Cas9 cells were washed with 1X phosphate-buffered saline, resuspended in 20 µL of solution SF (Lonza), and combined with 10 µM gRNA together with 0.5 µM of the GUIDE-seq dsODN. This mixture was transferred into one well of a 96-well Nucleocuvette plate (Lonza) and electroporated using protocol DS150. Following electroporation, cells were transferred to a 96-well plate preheated with Eagles Minimum Essential Medium (EMEM; ATCC: 30-2003). Cells were incubated at 37°C with 5% CO2 for 72 hours. After incubation, gDNA was extracted using QuickExtract (Biosearch Technologies). 250 ng of gDNA was fragmented, and P5 adapters were ligated using the xGen DNA Library Prep EZ UNI kit (IDT). GUIDE-seq library amplification, sequencing, and operation of the GUIDE-seq software were performed as previously described with the inclusion of a glocal implementation of Needleman-Wunch alignment for gRNA to target region alignment.

Comparison of GUIDE-Seq hits against TaP-N-Seq hits was performed in the same way as the *in silico* predictions.

### Endpoint and Live Cell Microscopy

Endpoint microscopy was captured on a Nikon Ti2 inverted microscope with a 20X/0.75NA Air Objective, Andor Zyla sCMOS camera, Lumencore SpectraX LED excitation, and Chroma ET filter cubes (DAPI 49000, GFP 49002, mCherry 49008, Cy5 49006). For multi-field stitched images in dim channels (DAPI, Cy5), background subtraction using shading correction within the Nikon Elements software. Cells were generally imaged in standard 12-well and 6-well culture dishes.

Live cell microscopy was captured on a Zeiss Axio Observer Z1 inverted microscope with a 10X/0.4NA Air Objective, Hamamatsu Orca R^2^ CCD camera, and Zeiss Colibri 2 LED excitation. An external Zeiss incubation system maintained the temperature, humidity, and CO^2^ environment. For live cell imaging, cells were cultured in ibidi TC-treated glass-bottom 24-well plates (ibidi Cat# 82427). Before starting a run, the system is allowed to equilibrate for 2 hours. Fields of view were selected, around which a 2x2 stitched pattern with 30% overlap was set. For each capture, the integrated Definite Focus determines the glass-liquid interface, after which images are captured based on pre-determined offsets for each channel. Post-transfection, cells were generally imaged for approximately 5 days at 2-hour intervals across all channels. After this, cells were passaged into a new plate at standard seeding concentration, allowed to expand briefly for approximately 2 days, then imaged for an additional 5 days.

Live cell images are then exported as 12-bit TIFF files for each scene, time, channel, and tile. These images are then imported into MATLAB. Each image tile is background corrected using shading correction (*imsubtract*), tiled, and contrasted. The processed images were then used to generate both a merged RGB image and a grayscale image for the channel of interest for each time point. Timestamps and scale bars were overlaid onto the images, then the image stack was exported as a .mp4 video.

### Statistical tests

Statistical tests are performed as one-tailed, two-sample student t-tests. Error bars shown are generally means and standard deviations, with the exception of flow cytometry fluorescence, for which geometric means and geometric standard deviations were used.

